# Basal autophagy during meiotic prophase I is required for accurate chromosome segregation in Drosophila oocytes and declines during oocyte aging

**DOI:** 10.1101/2025.05.13.653863

**Authors:** Diana C. Hilpert, Muhammad Abdul Haseeb, Sharon E. Bickel

**Author notes:** Department of Biochemistry and Cell Biology, Geisel School of Medicine at Dartmouth, 74 College St. Hanover, NH 03755.

## Abstract

Meiotic segregation errors in human oocytes are the leading cause of miscarriages and trisomic pregnancies and their frequency increases exponentially for women in their thirties. One factor that contributes to increased segregation errors in aging oocytes is premature loss of sister chromatid cohesion. However, the mechanisms underlying age-dependent deterioration of cohesion are not well-defined. Autophagy, a cellular degradation process critical for cellular homeostasis, is known to decline with age in various organisms and cell types. Here we quantify basal autophagy in Drosophila oocytes and use GAL4/UAS inducible knockdown to ask whether disruption of autophagy in prophase oocytes impacts the fidelity of chromosome segregation. We find that individual knockdown of autophagy proteins in Drosophila oocytes during meiotic prophase causes a significant increase in segregation errors. In addition, Atg8a knockdown in prophase oocytes leads to premature loss of arm cohesion and missegregation of recombinant homologs during meiosis I. Using an oocyte aging paradigm that we have previously described, we show that basal autophagy decreases significantly when Drosophila oocytes undergo aging. Our data support the model that a decline in autophagy during oocyte aging contributes to premature loss of meiotic cohesion and segregation errors.

## INTRODUCTION

Chromosome segregation errors during female meiosis, the leading cause of miscarriages and severe congenital syndromes in humans, increase dramatically as women age, a phenomenon known as the maternal age effect (Nagaoka *et al*., 2012; Wartosch *et al*., 2021). Analysis of over 20,000 human oocytes (based on polar body genotyping from IVF patients) revealed that 20% of the oocytes from 35-year old women gave rise to aneuploid eggs but this doubled to more than 40% for the oocytes from 40-year old women (Kuliev *et al*., 2011).

One factor that contributes to the maternal age effect is precocious loss of sister chromatid cohesion in aging oocytes (Duncan *et al*., 2012; Sakakibara *et al*., 2015; Zielinska *et al*., 2015; Webster and Schuh, 2017). Cohesion, mediated by cohesin complexes, keeps sister chromatids connected until their controlled segregation during the meiotic divisions. After meiotic exchange, cohesive linkages between sisters distal to the crossover keep recombinant homologs physically associated. In humans, meiotic recombination is completed in fetal oocytes that arrest before birth in an extended diplotene (dictyate) state in which the bivalent relies on arm cohesion to keep homologous chromosomes together until ovulation and anaphase I, which can happen decades later (Nagaoka *et al*., 2012). Normal segregation during the 2^nd^ meiotic division requires centromeric cohesion to remain intact until anaphase II. While it is clear that both arm and centromeric cohesion deteriorate as oocytes age (Subramanian and Bickel, 2008; Chiang *et al*., 2010; Duncan *et al*., 2012; Sakakibara *et al*., 2015; Zielinska *et al*., 2015; Webster and Schuh, 2017; Perkins *et al*., 2019), the mechanisms that lead to aging-induced loss of cohesion are not well-defined.

Autophagy is a cellular degradation process conserved from yeast to humans. Under nutrient deprivation, bulk autophagy ensures sufficient amino acid and energy supply. However, even under normal nutrient conditions, autophagy operates at a basal level that is required to maintain protein and organelle homeostasis (Aman *et al*., 2021). Basal levels of autophagy are known to decrease with age in various cell types and across species, including humans, contributing to a decline in cellular health in aging cells (Rubinsztein *et al*., 2011; Barbosa *et al*., 2018; Hansen *et al*., 2018; Lim *et al*., 2024).

Here we confirm that under normal feeding conditions, basal autophagy occurs in the Drosophila female germline and report that disruption of basal autophagy in Drosophila oocytes during meiotic prophase leads to premature loss of arm cohesion and meiotic chromosome segregation errors. In addition, we demonstrate that basal autophagy declines in Drosophila oocytes when they undergo aging. Our data support the model that an age-dependent decrease in autophagy contributes to deterioration of meiotic cohesion in Drosophila oocytes and increased chromosome segregation errors in aged oocytes.

## RESULTS AND DISCUSSION

### Basal autophagy occurs in the female germline under normal feeding conditions

The Drosophila ovary is an excellent model to study the effect of starvation on autophagy induction and cell death pathways (Pritchett *et al*., 2009; Lebo and McCall, 2021). Every ovary contains multiple ovarioles, each composed of an assembly line of egg chambers containing germ cells (one oocyte and 15 nurse cells), as well as somatic follicle cells that surround each egg chamber (**Figure S1B**). Previous studies have demonstrated that nutrient deprivation (starvation) induces autophagy in Drosophila ovaries (Hou *et al*., 2008; Barth *et al*., 2011). Furthermore, prolonged starvation can lead to induced programmed cell death that is accompanied and partially mediated by increased autophagy in the germarium and mid-stage egg chambers (Peterson *et al*., 2003; Hou *et al*., 2008; DeVorkin *et al*., 2014). While autophagy is required in follicle cells for egg chamber development, it is dispensable in germ cells for egg chamber development (Barth *et al*., 2011).

To characterize basal autophagy in the Drosophila female germline under normal feeding conditions, we performed immunostaining to detect autophagosomes and autolysosomes in the ovarioles of mated females. To label autophagic vesicles, we stained for Atg8a, a ubiquitin-like protein that is conjugated to autophagic membranes. To distinguish between autophagosomes and autolysosomes, we stained for Cathepsin L (CP1), a lysosomal marker. Whereas autophagosomes and autolysosomes are both positive for Atg8a, only autolysosomes are positive for CP1.

In three genotypes commonly used as “wild type”, we observed Atg8a foci in nurse cells, oocytes and follicle cells at multiple stages with large variance between and within genotypes (**Figure 1**). Absence of Atg8a foci in *atg7^d14^/ atg7^d77^* null trans-heterozygotes that lack Atg7, an E1-like activating enzyme required for the conjugation of Atg8a to membranes (Martens and Fracchiolla, 2020), confirms that the Atg8a foci detected are indeed autophagic vesicles (**Figures 1 & 2)**. For stage 7 oocytes (diplotene), the relative oocyte volume occupied by autophagosomes or autolysosomes varied 3-4 fold between the wild-type strains, with no obvious correlation between the volume of autophagosomes and autolysosomes. Both vesicle types were absent in *atg7* null oocytes (**Figures 1 & 2)**.

**Figure 1:**
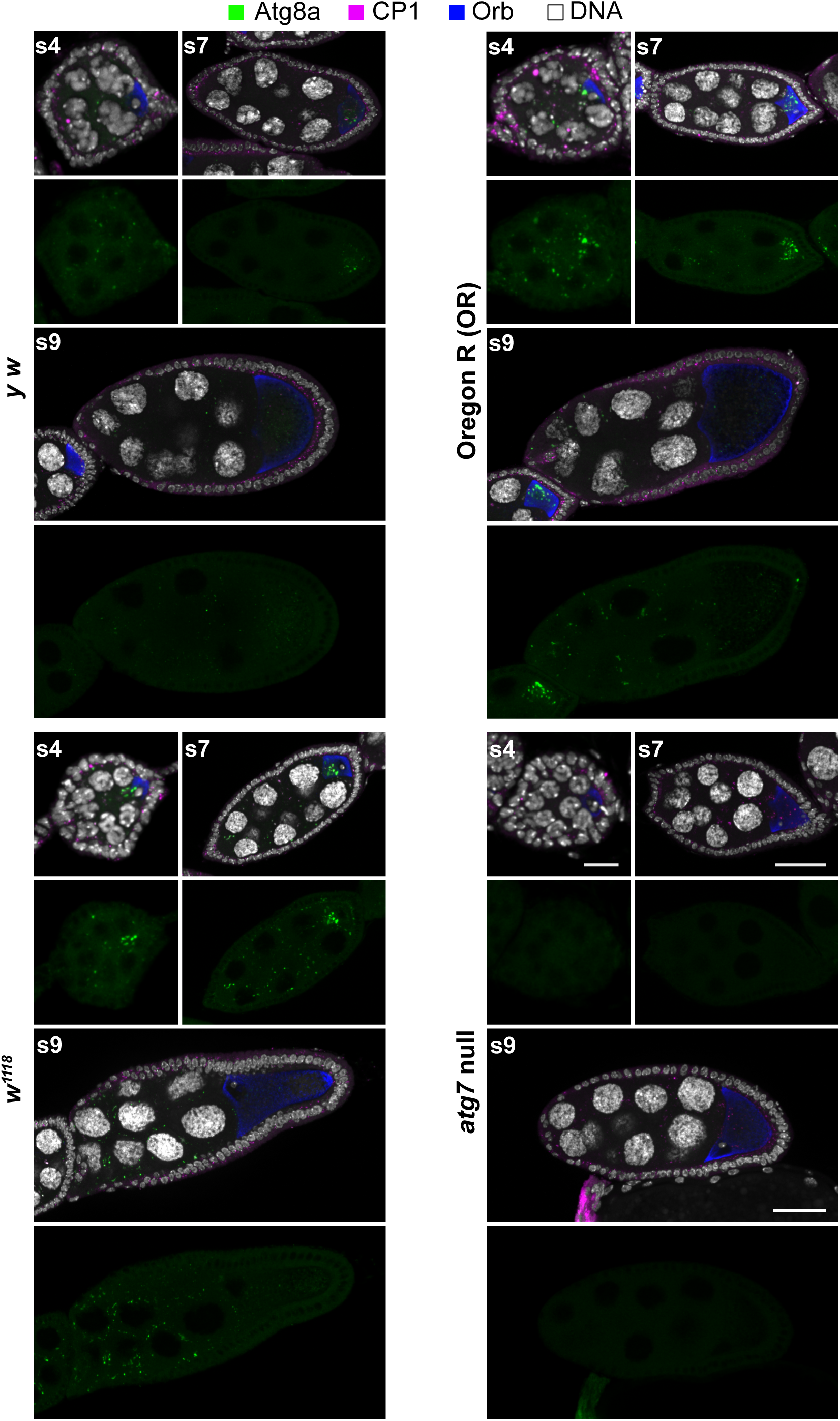
Basal autophagy occurs in the female germline throughout meiotic prophase. Images shown for stage 4 (s4), stage 7 (s7), and stage 9 (s9) egg chambers in three strains commonly utilized as “wild type” as well as *atg7* null females (*atg7^d14^/ atg7^d77^*). Atg8a staining (green) marks autophagic vesicles, Cathepsin L (CP1) signal (magenta) marks lysosomes, Orb (blue) is enriched in the oocyte cytoplasm, and DNA is shown in white. Atg8a foci are absent in *atg7* null egg chambers, validating the detection method for autophagic vesicles. Images are maximum intensity projections of confocal Z series. Scale bar for stage 4 is 5 µm and 30 µm for stages 7 and 9.

**Figure 2:**
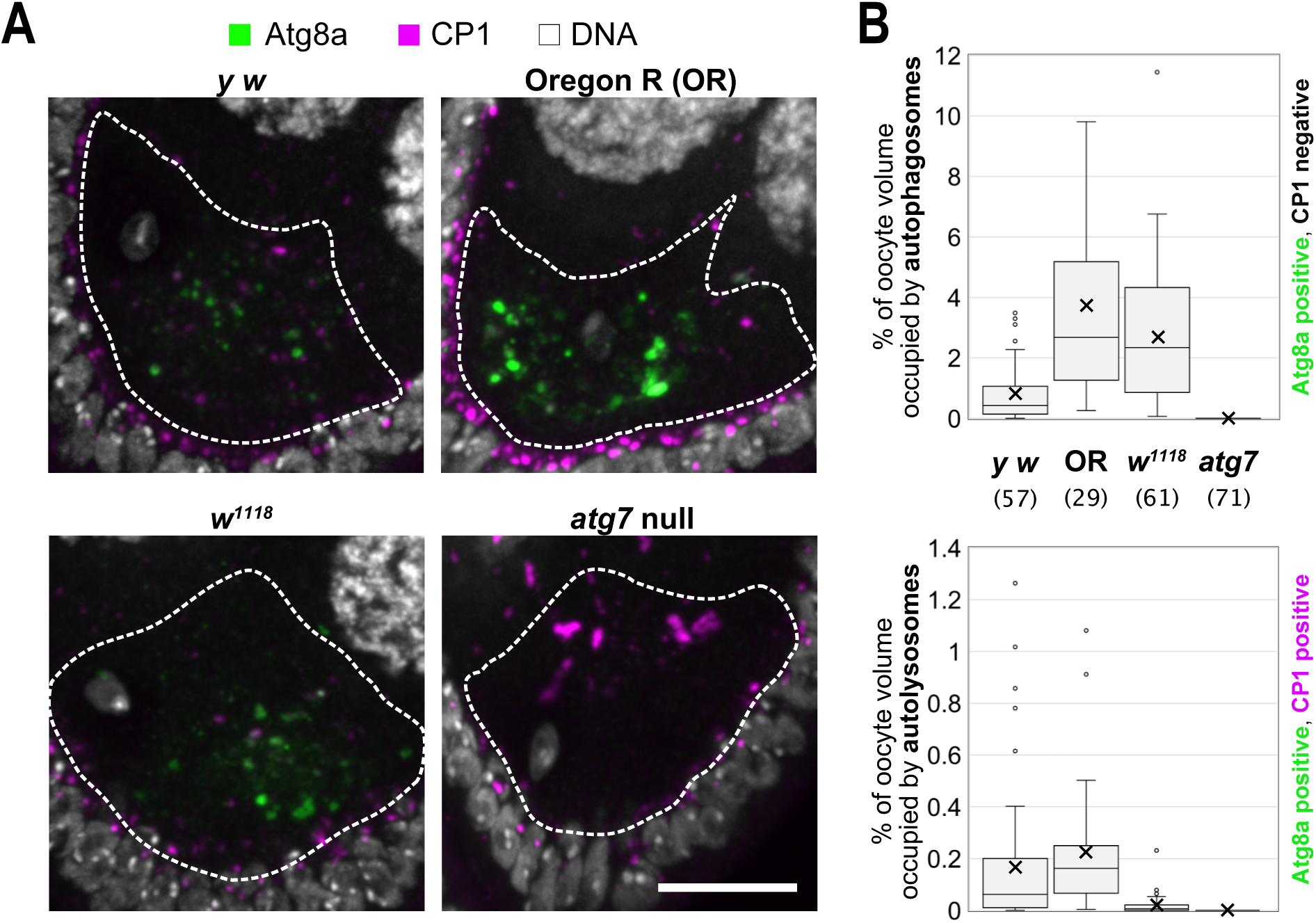
Quantification of basal autophagy in diplotene oocytes. **(A)** Autophagic vesicles in stage 7 (diplotene) oocytes. Hatched lines demarcate the oocyte, as determined by Orb signal (not shown). Atg8a (green), CP1 (magenta) and DNA (white). Foci that are Atg8a positive but lack CP1 signal are autophagosomes. Foci that are positive for both Atg8a and CP1 are autolysosomes. Maximum intensity projections of confocal Z series, scale bar, 10 µm. **(B)** Graphs present the percentage of oocyte volume occupied by autophagosomes (Atg8a positive, CP1 negative) and autolysosomes (Atg8a positive, CP1 positive). Number of oocytes quantified is indicated in parentheses for each genotype.

We also used GAL4/UAS induction **(Figure S1A)** to knock down Atg3, an E2-like ubiquitin ligase that is also required for the Atg8a conjugation to membranes (Fang *et al*., 2021). Use of the maternal alpha tubulin GAL4-VP16 driver (hereafter termed matα driver) allows us to induce UAS-hairpin expression in the germline after meiotic S phase with levels that increase during prophase I (**Figure S1B).** In Atg3 KD oocytes, we observed a significant reduction in the oocyte volume occupied by autophagosomes and autolysosomes, as expected **(Figure S1 C&D)**, further confirming that our immunodetection technique is specific for autophagic vesicles and that basal autophagy significantly decreases when essential components of the conjugation machinery are reduced. Our data demonstrate that in the absence of starvation, basal autophagy is detectable within the Drosophila female germline of multiple wild-type strains, but levels and location (oocyte and/or nurse cells) vary widely depending on the genotype and egg chamber stage. Even though absolute levels of basal autophagy vary across genotypes and are quite low in Atg3 KD and control genotypes, a significant decrease in autophagic vesicles is clearly detectable when Atg3 is knocked down during meiotic prophase.

### Autophagy is required during prophase I for accurate meiotic chromosome segregation in Drosophila oocytes

To ask whether basal autophagy is required for accurate segregation in Drosophila oocytes under normal feeding conditions, we used the matα driver to knock down Atg8a during meiotic prophase (**Figure S2A**, **Figure 3A**) and performed a genetic assay **(Figure S2C)** to measure X chromosome nondisjunction (NDJ) in control and KD oocytes. We use the term NDJ to broadly describe any type of segregation error. Females were also heterozygous for the *mtrm^KG^* allele **(Figure S2A)** which disables the achiasmate segregation pathway in Drosophila oocytes so that premature loss of arm cohesion results in missegregation **(Figure S2B)**.

**Figure 3:**
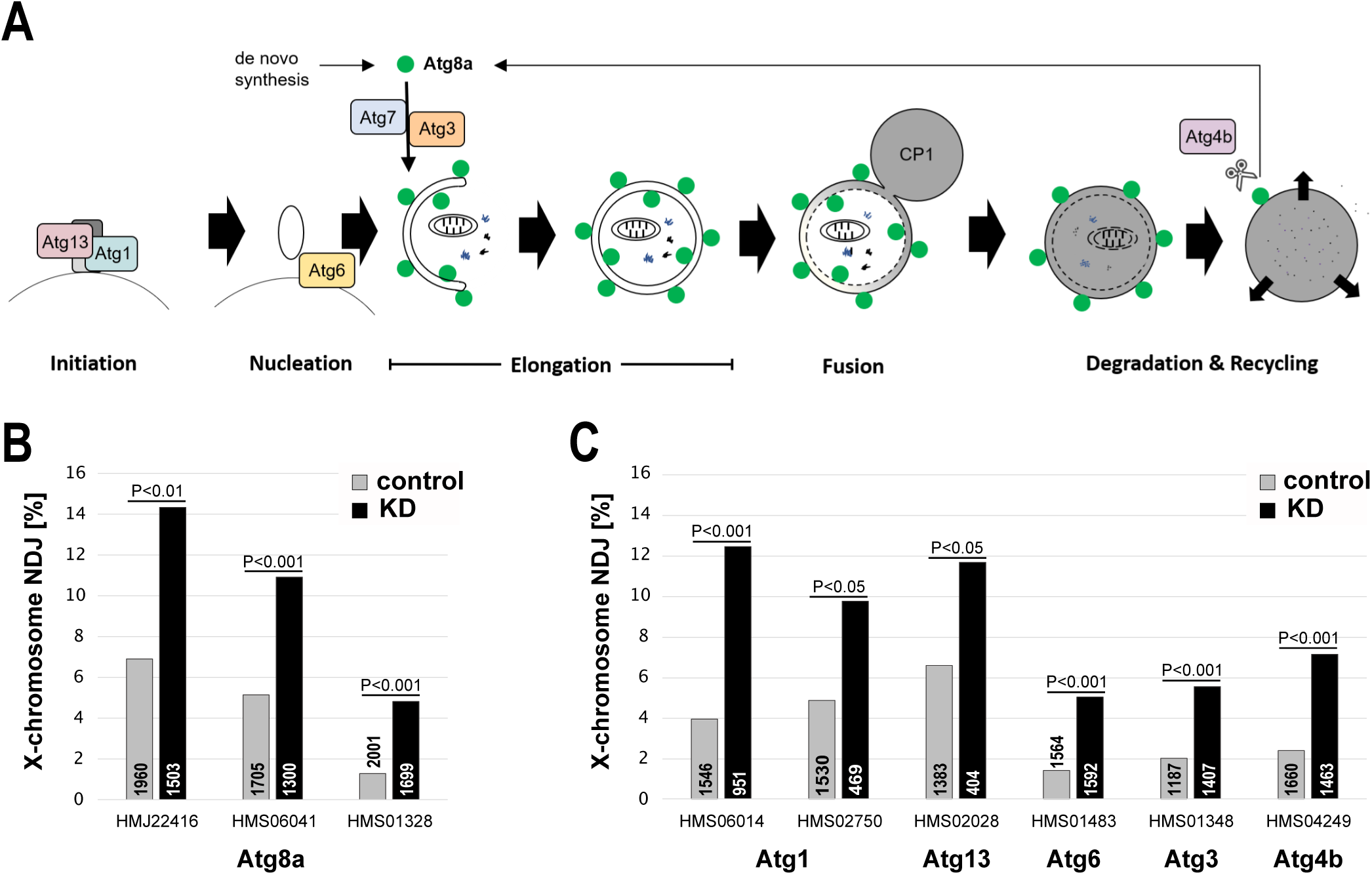
Germline knockdown of different autophagy proteins during meiotic prophase significantly increases oocyte segregation errors. **(A)** Cartoon illustrates steps in the autophagy pathway with proteins targeted for knockdown shown in colored boxes. (**B and C)** For each protein shown, the matα driver was used to induce knockdown during meiotic prophase. X-chromosome segregation errors (NDJ) were quantified in knockdown (KD) and control oocytes. Total number of progeny scored for each genotype is presented within or directly above the bars and the hairpin identifier is shown for each pair. All genotypes tested were heterozygous for *mtrm^KG^* to disable the achiasmate segregation system that operates in Drosophila oocytes. When functional, this system would ensure accurate segregation of bivalents that lose their chiasmata due to premature loss of arm cohesion (see **Figure S2B**).

Knockdown of Atg8a, which decreases autophagosome formation (Nakatogawa *et al*., 2007; Xie *et al*., 2008; Martens and Fracchiolla, 2020), caused a significant increase in meiotic NDJ (**Figure 3B**). For three different Atg8a hairpins, we observed that segregation errors were significantly higher in matα→Atg8a KD oocytes than their corresponding controls.

We also quantified X-chromosome NDJ in oocytes in which we induced individual knockdown of additional proteins required for autophagy (Kitada and Koya, 2021; Lim *et al*., 2024) (**Figure 3A, Figure S2A)**. Prophase I specific knockdown of proteins involved in autophagy initiation (Atg1, Atg13), phagophore nucleation (Atg6), Atg8a conjugation (Atg3) or recycling (Atg4b) each caused a significant increase in meiotic chromosome segregation errors (**Figure 3A&C)**. Importantly, whereas the Atg8a conjugation system is also required for LC3- associated endocytosis (LANDO) and LC3-associated phagocytosis (LAP), the autophagy initiation complex comprised of Atg1 and Atg13 is dispensable for these pathways (Peña-Martinez *et al*., 2022). Together, these data demonstrate that accurate chromosome segregation in Drosophila oocytes depends on basal autophagy during meiotic prophase.

### Cohesion defects increase significantly when autophagy is disrupted during prophase I

To ask whether basal autophagy is required for cohesion maintenance in Drosophila oocytes during meiotic prophase, we quantified X chromosome arm and centromeric cohesion in Atg8a KD and control oocytes using Fluorescence In-Situ Hybridization (FISH). We assayed arm cohesion using an Oligopaint probe that hybridizes to a 100-kb region on the distal X chromosome arm. To score for centromeric cohesion defects we used a 359-bp satellite repeat probe that hybridizes to an 11MB region near the centromere of the X-chromosome (**Figure 4A**). For a given probe, one or two spots within the bivalent were quantified as intact cohesion and three or four spots were counted as premature loss of cohesion (**Figure 4B**). Using these probes and FISH assay, we have previously demonstrated that matα-induced knockdown of the cohesin subunit SMC3 or the cohesin loader Nipped-B causes a significant increase in the percentage of oocytes with arm cohesion defects, validating this method (Haseeb *et al*., 2024a; Haseeb *et al*., 2024b).

**Figure 4:**
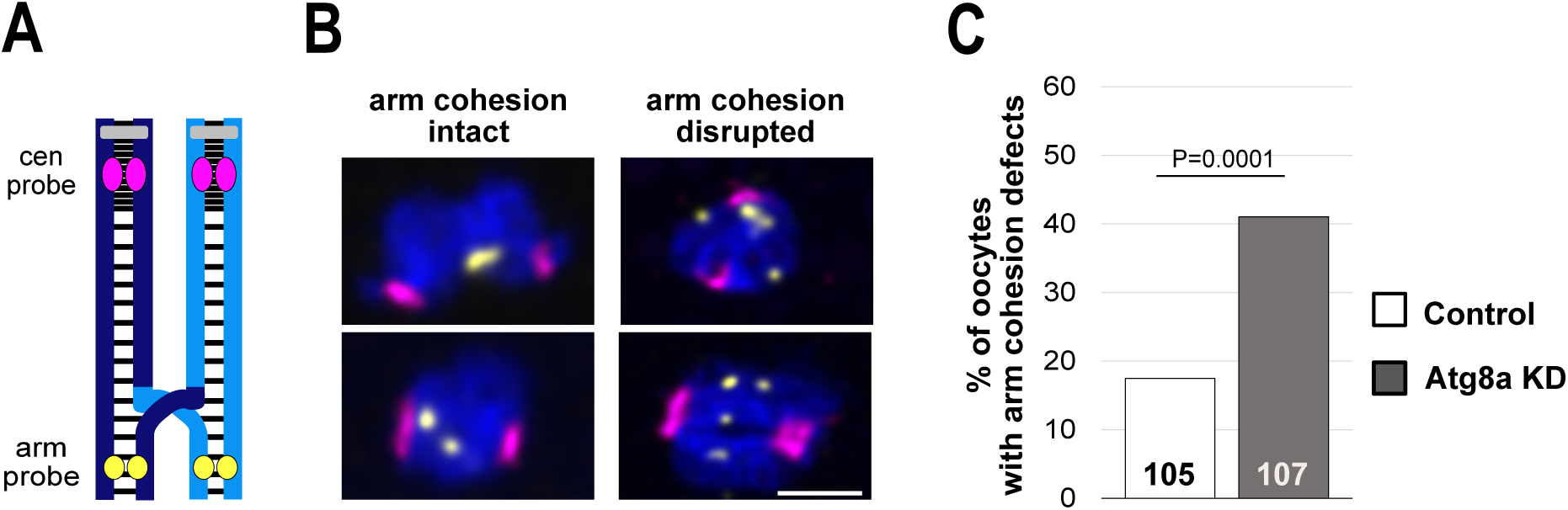
Oocytes with arm cohesion defects increase when Atg8a is knocked down during meiotic prophase. **(A)** Drosophila X chromosome bivalent with a single crossover is shown with centromeres in gray and cohesion represented as black lines. Sites of hybridization for the arm probe (yellow) and centromere proximal probe (magenta) are illustrated. **(B)** Examples of intact arm cohesion (1 or 2 yellow spots) and premature loss of arm cohesion (3 or 4 yellow spots). Maximum intensity projections of confocal Z series, scale bar, 2 μm. **(C)** Percentage of oocytes with arm cohesion defects in Atg8a KD (matα driver → *Atg8a^HMS06041^* hairpin) and control (matα driver → *mCherry^TB00168^*hairpin). The number of oocytes scored is shown in each bar.

We compared cohesion in matα→ Atg8a KD oocytes and matα→ mCherry hairpin oocytes (same driver, no target) and found that arm cohesion defects were significantly elevated in Atg8a KD oocytes (**Figure 4C**). In contrast, we did not detect any centromeric cohesion defects, possibly due to the large size of satellite DNA near the centromere. Because deterioration of arm cohesion can lead to chiasma destabilization and random segregation of recombinant homologs during anaphase I, we performed an additional genetic assay **(Figure S2D)** to determine whether recombinant homologs missegregate more frequently when Atg8a is knocked down. Consistent with precocious loss of arm cohesion, Atg8a knockdown caused a significant increase in the frequency at which recombinant homologs (but not sisters) missegregate (**Figure S2E**). However, Atg8a KD does not change the location or frequency of crossovers along the X chromosome **(Figure S2F),** so increased missegregation of recombinant homologs in Atg8a KD oocytes is not due to increased meiotic exchange. Our data indicate that under normal feeding conditions, maintenance of arm cohesion in Drosophila oocytes depends on basal autophagy during meiotic prophase.

### Autophagy declines in diplotene oocytes that undergo aging

Oogenesis in Drosophila is normally continuous, and prophase oocytes do not arrest and age. However, our lab has developed a standardized protocol to arrest and age diplotene oocytes within the Drosophila ovary (Subramanian and Bickel, 2008). In the absence of mating, egg laying is suppressed in females, and most oocyte stages halt development and age within the ovary. After mating, aged oocytes mature, are fertilized and can be scored for segregation errors. In our 4-day aging regimen, diplotene is extended 11-18 fold for oocytes that undergo aging (Subramanian and Bickel, 2008).

Our aging experiments utilize a sensitized genetic background in which females are heterozygous for *smc1Δ* and *mtrm^KG^* mutations. In *smc1Δ*/+ oocytes, the number of functional cohesin complexes is reduced so four days of aging is sufficient to cause premature loss of cohesion. Heterozygosity for *mtrm^KG^* disables the achiasmate pathway so that recombinant homologs that lose cohesion will segregate randomly **(Figure S2B)**. Recent work also indicates that lowering the dosage of Mtrm weakens arm cohesion during meiotic prophase (Bonner *et al*., 2020; Haseeb *et al*., 2024b). We have previously demonstrated that when diplotene *mtrm^KG^/smc1Δ* oocytes undergo aging, NDJ is significantly increased compared to oocytes that do not age (Subramanian and Bickel, 2008; Perkins *et al*., 2019).

To determine whether basal autophagy is reduced during oocyte aging under normal feeding conditions, we slightly modified our aging regimen so that it utilizes the same cornmeal-molasses food used for our other experiments **(Figure S3)**. We confirmed that virgin *mtrm smc1Δ/+* females held their eggs **(Figure S3)** and that NDJ was significantly higher in aged oocytes than non-aged oocytes (**Figure 5A**). We quantified basal autophagy in stage 7 (diplotene) oocytes that did and did not undergo aging (**Figure 5B**) and found that aging caused a significant decline in the oocyte volume occupied by autophagosomes and autolysosomes (**Figure 5C**).

**Figure 5:**
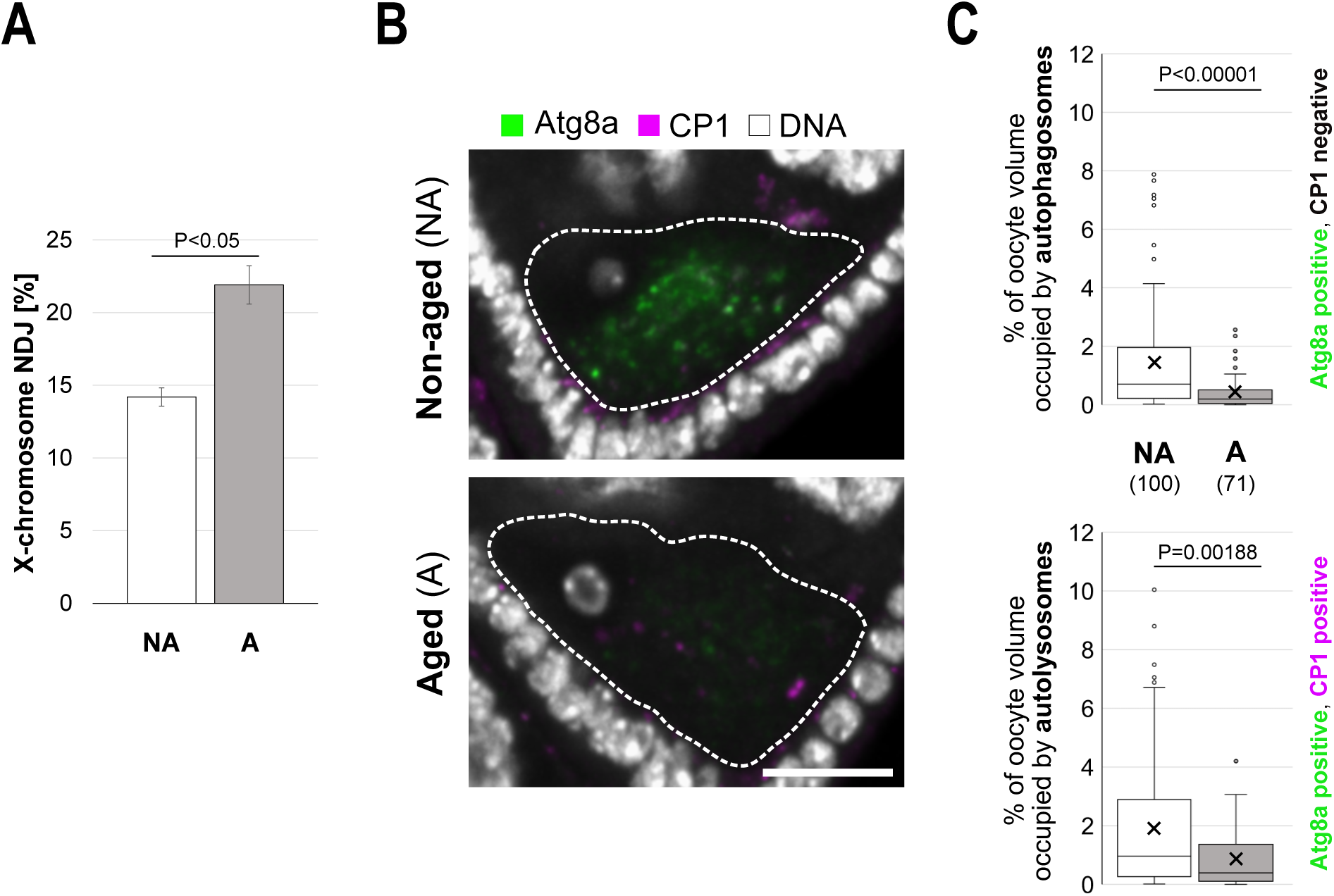
Basal autophagy declines in diplotene oocytes that undergo aging. **(A)** X-chromosome NDJ was measured for aged (A) and non-aged (NA) oocytes after subjecting *mtrm^KG^ smc1Δ/+* mothers to an aging regimen that utilized standard cornmeal- molasses food. The average of four tests is graphed, error bars depict SEM, and P value is < 0.05 for each individual test. **(B)** Images of aged and non-aged stage 7 (diplotene) oocytes from *mtrm^KG^ smc1Δ/+* mothers. Atg8a (green), CP1 (magenta) and DNA (white). Hatched lines outline the perimeter of the oocyte, as determined by Orb signal (not shown). Maximum intensity projections of confocal Z series, scale bar, 10 μm. **(C)** Quantification of basal autophagy in stage 7 aged (A) and non-aged (NA) oocytes of *mtrm^KG^ smc1Δ/+* females. The percentage of oocyte volume occupied by autophagosomes (Atg8a positive, CP1 negative) and autolysosomes (Atg8a positive, CP1 positive) is significantly decreased when oocytes undergo aging. The number of oocytes scored is shown in parentheses.

Our study demonstrates that basal autophagy during meiotic prophase is required for cohesion maintenance and accurate chromosome segregation in Drosophila oocytes.

Furthermore, we find that autophagy declines when diplotene oocytes undergo aging. These data support the model that a decrease in basal autophagy during oocyte aging contributes to the maternal age effect in humans. Because stimulation of autophagy in aging cells has been shown to improve cell function (Gelino and Hansen, 2012; Aman *et al*., 2021; Li *et al*., 2021), therapeutic approaches to preserve or increase autophagy in aging oocytes may provide an effective strategy to reduce meiotic segregation errors in older women.

## MATERIALS AND METHODS

### Fly stocks and crosses

Stocks and crosses were raised on standard cornmeal-molasses food and kept at 25°C in a humidified incubator. Table S1 provides full genotypes for all stocks used in this study as well as Bickel stock numbers which are mentioned below for cross schemes.

### Aging regimen

We typically utilize laying bottles with glucose agar plates and a smear of wet yeast paste on each plate for our oocyte aging regimen (Subramanian and Bickel, 2008; Perkins *et al*., 2019). However, to investigate the effects of oocyte aging on basal autophagy in the female germline, we wanted to keep the food constant throughout the experiment. Therefore, we altered our aging regimen to utilize the same standard cornmeal-molasses fly food that we consistently use for fly rearing and NDJ tests **(see Figure S3)**.

*y w* virgin females (A-062) were crossed to *mtrm^KG^ smc1Δ /TM3* males (M-822) to generate female progeny (*mtrm^KG^ smc1Δ / +*) that were collected as virgin females within an 8- 12 hour window and placed overnight in standard cornmeal-molasses food vials with a sprinkle of dry yeast. Aging regimen bottles were set up at approximately 2pm and flies were transferred to a new bottle (by flipping, no CO2) every 24 hours over a 4-day interval. To generate non-aged oocytes, 45 virgins were placed in a bottle with 20 *X^Y, Bar* (C-200). Because oogenesis and egg laying occur in an uninterrupted manner when females are mated, their oocytes do not arrest or undergo aging. To generate aged oocytes, 45 virgins were placed into a bottle without males. In the absence of mating, egg laying is suppressed and oogenesis halts for oocytes at stages 8 and earlier (Subramanian and Bickel, 2008). These oocytes arrest and age within the female ovary.

Each time flies were transferred, images were captured of the food surface of each bottle to confirm egg laying or holding. After the 4-day aging regimen, females were immediately processed for NDJ tests or immunostaining. We limit the aging regimen to 4 days because egg laying increases for unmated females after this time period.

### X-chromosome NDJ test

To measure chromosome segregation errors (NDJ) in control and KD oocytes, *y w; + ; mtrm^KG^/TM3, Sb* (W-109) or *y w; + ; mtrm^KG^, matα/TM3, Sb* (W-110) males were crossed to virgins containing a UAS-hairpin transgene. Non-balancer **control** (mtrm^KG^, no driver → UAS- hairpin) and **KD** (mtrm^KG^, matα driver → UAS-hairpin) virgin female progeny **(Figure S2A)** were collected over a 5-day period and held in vials containing standard cornmeal-molasses food with a sprinkle of dry yeast. For each NDJ test, control and KD virgins were crossed to *X^Y, Bar* (C- 200) males. For age-dependent NDJ experiments, *mtrm^KG^ smc1Δ / +* females were crossed to *X^Y, Bar* (C-200) males after completion of the aging regimen.

For each genotype, 10 vials were started (day 0) with 4-6 females and 2-3 males per vial (standard food + dry yeast sprinkle). The parents were removed on day 7 and progeny were scored through day 18. Because only half of the progeny arising from exceptional (aneuploid) gametes survive but all the progeny arising from normal gametes survive (Figure S2B), % NDJ is calculated using the following formula: [(2(Diplo-X + Nullo-X))/(N + 2(Diplo-X + Nullo-X))]100, where N is the total number of progeny scored. The calculator provided by Gilliland and colleagues (Zeng *et al*., 2010) was used to determine whether NDJ was significantly different between control and KD genotypes for each hairpin.

### Recombinational history assay

To determine whether matα → Atg8a KD alters the frequency at which recombinant homologs and/or sisters missegregate, *y sc cv v f car; + ; mtrm^KG^/TM3, Sb* (M-835) or *y sc cv v f car; + ; mtrm^K^, matα/TM3, Sb* (M-834) males were crossed to *y* virgins containing the UAS- *Atg8a^HMS06041^*hairpin insertion (I-585). NDJ was measured by crossing *y sc cv v f car / + ; Atg8a*^HMS06041^ */ +; mtrm^KG^/ +* **(control)** and *y sc cv v f car / + ; Atg8a*^HMS06041^ */ +; mtrm^KG^ matα/ +* **(KD)** virgins to *X^Y, Bar* (C-200) males. NDJ tests were scored daily through day 18, and each Diplo-X female (arising from missegregation) was phenotyped for *s, cv, f,* and *car* and crossed to two *y w* (A-062) males. By scoring her male progeny for *sc, cv, f,* and *car*, one may deduce the X chromosome genotype of each Diplo-X female to determine whether either of the X chromosomes she inherited recombined prior to missegregation and whether homologs (*car ^+/-^*) or sisters (*car^+/+^* or *car^-/-^*) missegregated.

The frequency at which recombinant X chromosomes underwent missegregation in each genotype was calculated by dividing the number of Diplo-X females that inherited at least one recombinant chromosome by the total number of progeny scored in the NDJ test and multiplying by 100. The inheritance of two homologs (meiosis I) or two sisters (meiosis II) was graphed individually. A two-tailed, 2 x 2 *chi^2^* contingency test with Yates’ correction (GraphPad) was used to determine whether Atg8a KD significantly impacted the frequency at which Diplo-X females inherited at least one recombinant chromosome. P-values were determined separately for sisters and homologs.

Not all Diplo-X females yielded sufficient male offspring to confidently determine her genotype. In addition, although *car* is closely linked to the centromere (3.5cM), a rare crossover is possible. Overall, this assay provides a conservative frequency because the number of recombinant bivalents that missegregate will be underestimated given that double crossovers within the large *cv-f* interval are invisible and only two of the four chromatids in the bivalent will be inherited by the Diplo-X female and therefore be scored.

### X-chromosome crossover analysis

To ask whether matα → Atg8a KD alters frequency and/or placement of X chromosome crossovers, the same UAS-*Atg8a*^HMS06041^ control and KD virgin genotypes used above were crossed to *y w* (A-062) males and male progeny were scored for *sc*, *cv*, *f* and *car*. Fisher’s exact tests were performed in GraphPad to determine significance.

### Immunostaining

To visualize and quantify autophagic vesicles in oocytes, 10-12 females were held with 5-6 males in a vial containing standard cornmeal-molasses food and dry yeast for 1-3 days before dissection. *y w* (A-062) and *matα* (T-273) males were crossed to *Atg3^HMS01348^*virgins (I- 597) to generate control and KD egg chambers for immunostaining. To measure the effect of oocyte aging on basal autophagy, *mtrm^KG^ smc1Δ/+* females were subjected to the 4-day aging regimen (as described above) and ovaries from mated (non-aged oocytes) and non-mated (aged oocytes) were immediately dissected at the conclusion of the aging regimen.

We adapted the ovary immunostaining procedure developed by Shravage and colleagues (Nilangekar *et al*., 2019). All incubations and washes were performed in a covered glass dish with gentle agitation on an orbital shaker at room temperature unless noted otherwise. Ovaries were dissected in Grace’s media and ovarioles partially separated at their anterior end with the use of a tungsten needle (⁤ 15 minutes) followed by fixation for 15 minutes in 4% EM grade formaldehyde diluted in PBS and one brief rinse in PBSTx. After three 5-minute washes in PBSTx (PBS containing 0.1% Triton X), ovaries were permeabilized and blocked for one hour at room temperature in PBS containing 1% Triton X and 0.5% Bovine Serum Albumin (BSA). Ovaries were incubated with rabbit anti-Atg8a (1:100, ab109364, Abcam), guinea pig anti-Cathepsin L (1:200, Kinser and Dolph (2012)), and mouse monoclonal anti-Orb (1:30 each, 4H8 and 6H4, DSHB) in PBS containing 0.3% Triton X and 0.5% BSA with gentle rotation overnight at 4°C. The next day, samples were rinsed once and washed once in PBSTx for 15 minutes and blocked in PBSTx containing 10% Normal Donkey Serum (NDS) for two hours. Blocking was followed by a 2-hour incubation in secondary antibodies (Cy3 Donkey anti-rabbit, Cy5 Donkey anti-guinea pig and Alexa488 Donkey anti-mouse) diluted 1:400 in PBSTx containing NDS and protected from light. Following three 15-minute washes in PBSTx, ovaries were stained with DAPI (1 µg/mL in PBSTx) for 30 minutes followed by three 5-minute washes in PBSTx. Tungsten needles were used to separate ovarioles before mounting onto poly-L- lysine coated covereslips (18mm, #1.5) with Slowfade Glass.

### Fluorescence In-Situ Hybridization (FISH) to detect cohesion defects

To generate Atg8a KD and control genotypes for FISH analysis of cohesion defects, matα (T-273) males were crossed to virgins containing the Atg8a^HMS06041^ hairpin (H-137) or the mCherry^TB00168^ hairpin (B-042). 25 young female progeny from each cross were held with 10 males in a vial with standard cornmeal-molasses fly food and dry yeast for 3 - 4 days before dissection.

Fixation, predenaturation, hybridization, washes and mounting were performed as described previously (Haseeb *et al*., 2024b). To detect arm cohesion defects in prometaphase and metaphase I oocytes (stages 13 – 14), we utilized an Alexa 647-labeled Oligopaint probe (OPP122) generated by the Joyce Lab, University of Pennsylvania. This arm probe contains a mixture of 80 bp oligonucleotides that target a 100kb distal region of the X chromosome. To detect pericentric cohesion defects, we used a Cy3-conjugated probe (5’-Cy3- AGGGATCGTTAGCACTCGTAAT; Integrated DNA Technologies) that hybridizes to the satellite DNA repeats near the centromere of the X chromosome. Due to the large size of this region (11Mb), detection of centromeric cohesion defects are likely hampered.

### Image acquisition, processing and quantification

All imaging was performed using an Andor spinning disc confocal (50 µm pinhole) running Nikon Elements (5.11.02 Build 1369) to operate a Nikon Eclipse Ti inverted microscope, with an ASI MS 2000 motorized piezo stage, Zyla 4.2-megapixel sCMOS camera and four lasers (405, 561, 531 and 637 nm). 4X frame averaging (no binning) was used for all image acquisition, and Z series were captured by acquiring a complete Z stack for each fluor, progressing from the longest to shortest wavelength.

#### Immunostaining imaging

For egg-chamber imaging (Figure 1), a Nikon CFI 40X Plan Apo oil objective (NA 1.4) was utilized to collect a 5 µm Z series (0.2µm steps). For quantification of autophagosomes and autolysosomes in diplotene oocytes, a Nikon CFI 60X Plan Apo oil objective (NA 1.4) was used to capture a Z series (0.2 µm step size) that included the entire oocyte. Laser intensity and exposure time were adjusted for each experiment but kept constant for all groups within an experiment. For each figure presented, all image processing was identical for Atg8a signal in all panels and separately for CP1 signal in all panels. Representative images were chosen for each figure and the same number of optical sections are included within the maximum intensity projections presented for each panel in the figure.

#### FISH imaging

A CFI 100x oil Plan Apo DIC objective (NA 1.45) was used to capture a 4 µm Z series (512 x 512 pixels) with optical sections acquired every 0.1 µm.

#### Quantification of autophagic vesicles

The oocyte volume occupied by autophagosomes and autolysosomes was quantified using the Volocity 6.5.0 Quantification Module (Quorum Technologies). Orb is an RNA-binding protein that is enriched in the oocyte cytoplasm (Lantz *et al*., 1994). We utilized Orb signal intensity thresholding to identify all voxels corresponding to the oocyte volume. In some cases, manual cropping was needed to remove follicle cells from the volume and in these cases the full oocyte volume was not included in the analysis. Atg8a signal thresholding was used to identify autophagic vesicles and Cathepsin L signal thresholding was used to identify lysosomes. Signal thresholds to identify Atg8a and CP1 foci within the Orb volume were chosen conservatively so our quantification most likely underestimates the volume occupied by specific types of foci.

Quantification of autolysosomes was accomplished by identifying voxels occupied by Atg8a and CP1 signals that satisfied each threshold. To calculate the autophagosome volume in the oocyte, we subtracted the total volume corresponding to autolysosomes from the total volume occupied by Atg8a foci. Thresholds for Atg8a and CP1 were kept constant for all groups within an experiment and were slightly adjusted for each experiment based on cytosolic background.

Because the values for autophagosome and autolysosome volumes were not normally distributed (Shapiro-Wilk test), a non-parametric Mann-Whitney *U* test was utilized to test whether differences were statistically significant between groups (P value < 0.05). For all box and whiskers plots, an “X” marks the average intensity, with the inclusive median and quartiles depicted by horizontal lines and outliers shown as dots. Outliers correspond to values that lie more than 1.5 * IQR (interquartile range) above the 75% or below the 25% quartile.

#### Scoring cohesion defects

A MATLAB script was used to randomize and blind the FISH images before scoring. Image stacks were deconvolved (Volocity Restoration, v6.5.0) using identical parameters for both genotypes. Cohesion defects were scored manually by scrolling through the image stack and visualizing each spot in XY, XZ and YZ space. Two foci were scored as a “cohesion defect” only if there was no visible thread connecting them and the distance separating them was greater than ½ the diameter of the smallest spot. A two-tailed Fisher’s exact test (Graphpad) was utilized to determine whether the percentage of oocytes with arm defects were significantly different in Atg8a KD and control. No pericentromeric cohesion defects were detected in either genotype.

## ACKNOWLEDGEMENTS

We thank the Bloomington Drosophila Stock Center (NIH P40OD018537), the Transgenic RNAi Project (NIH R24OD030002) and Dr. Andreas Jenny for providing flies. 4H8 and 6H4 mouse anti-Orb monoclonal antibodies developed by Paul Schedl were obtained from the Developmental Studies Hybridoma Bank, created by the NICHD of the NIH and maintained by the Department of Biology at The University of Iowa. We are grateful to Patrick Dolph for the Cathepsin L (CP1) antibody. All Microscopy was performed in the Dartmouth Life Sciences Light Microscopy Core Facility. We thank Ann Lavanway for microscopy assistance and Britton Johnson for fly food preparation. We are grateful to Christopher Shoemaker for comments on the manuscript. DCH was funded, in part, by the Charles Gilman, ’52 Graduate Research Fellowship at Dartmouth. This work was funded by NIH R01GM059354 awarded to SEB.

---

**Figure S1:**
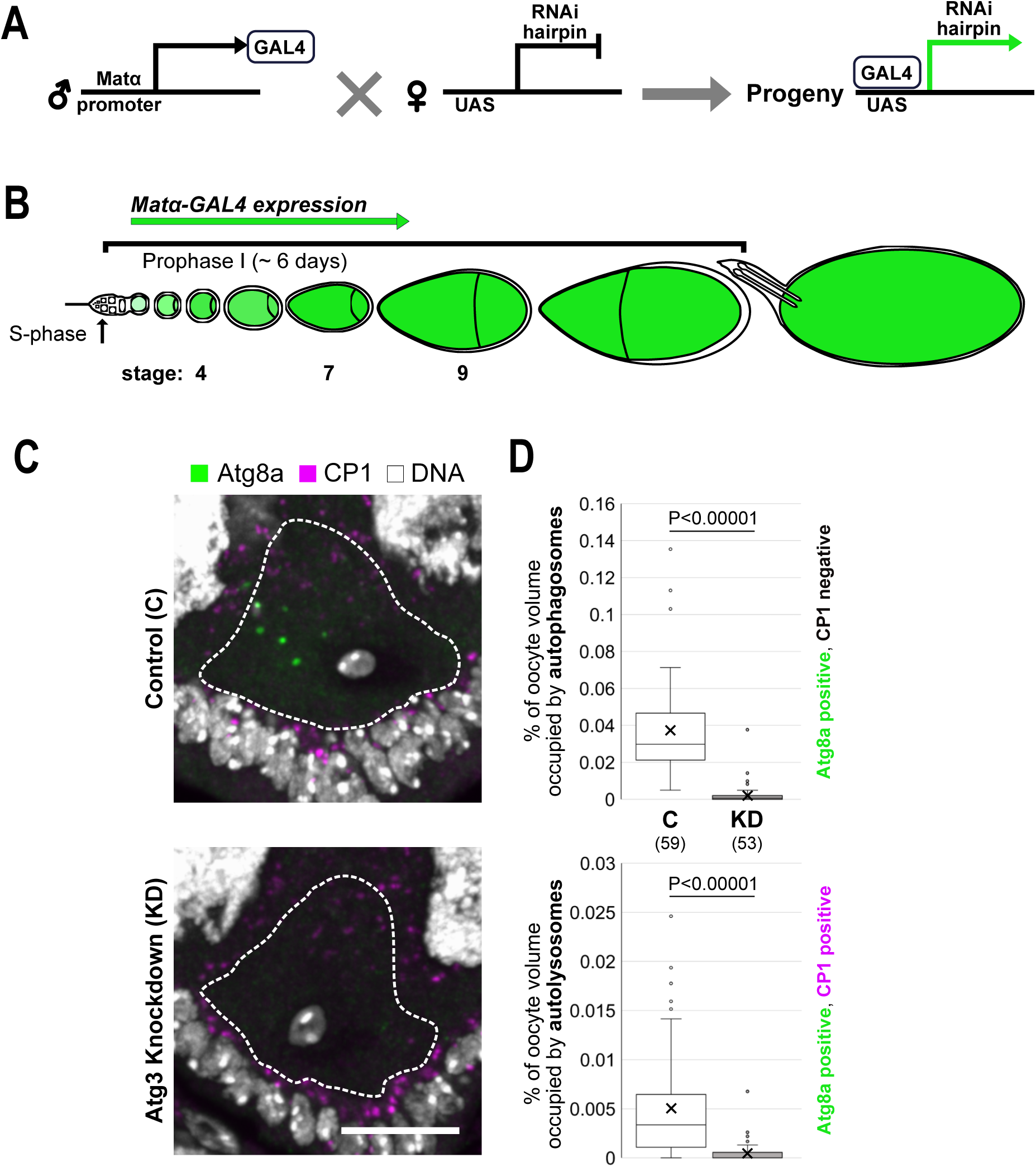
Atg3 knockdown during meiotic prophase decreases basal autophagy. **(A)** UAS/GAL4 method to induce RNAi hairpin expression and knock down proteins in the Drosophila female germline. **(B)** Diagram of Drosophila ovariole shows expression of the matα- GAL4 driver which begins after meiotic S-phase, when cohesion is first established. Because driver expression is restricted to prophase I, we can assay specifically for the effect of knockdown on cohesion maintenance. **(C)** Representative images of stage 7 (diplotene) oocytes are shown for Atg3 control (no driver → *Atg3^HMS01348^* hairpin) and Atg3 KD (matα driver → *Atg3^HMS01348^* hairpin). Hatched lines surround the oocyte, as determined by Orb signal (not shown). Autophagic vesicles are marked by Atg8a (green), lysosomes are marked by Cathepsin L (CP1, magenta) and DNA is shown in white. Maximum intensity projections of confocal Z series, scale bar, 10 µm. **(D)** Graphs that quantify basal autophagy in Atg3 KD and control oocytes (stage 7, diplotene) present the percentage of oocyte volume occupied by autophagosomes (Atg8a positive, CP1 negative) and autolysosomes (Atg8a positive, CP1 positive). Number of oocytes analyzed is shown in parentheses.

**Figure S2:**
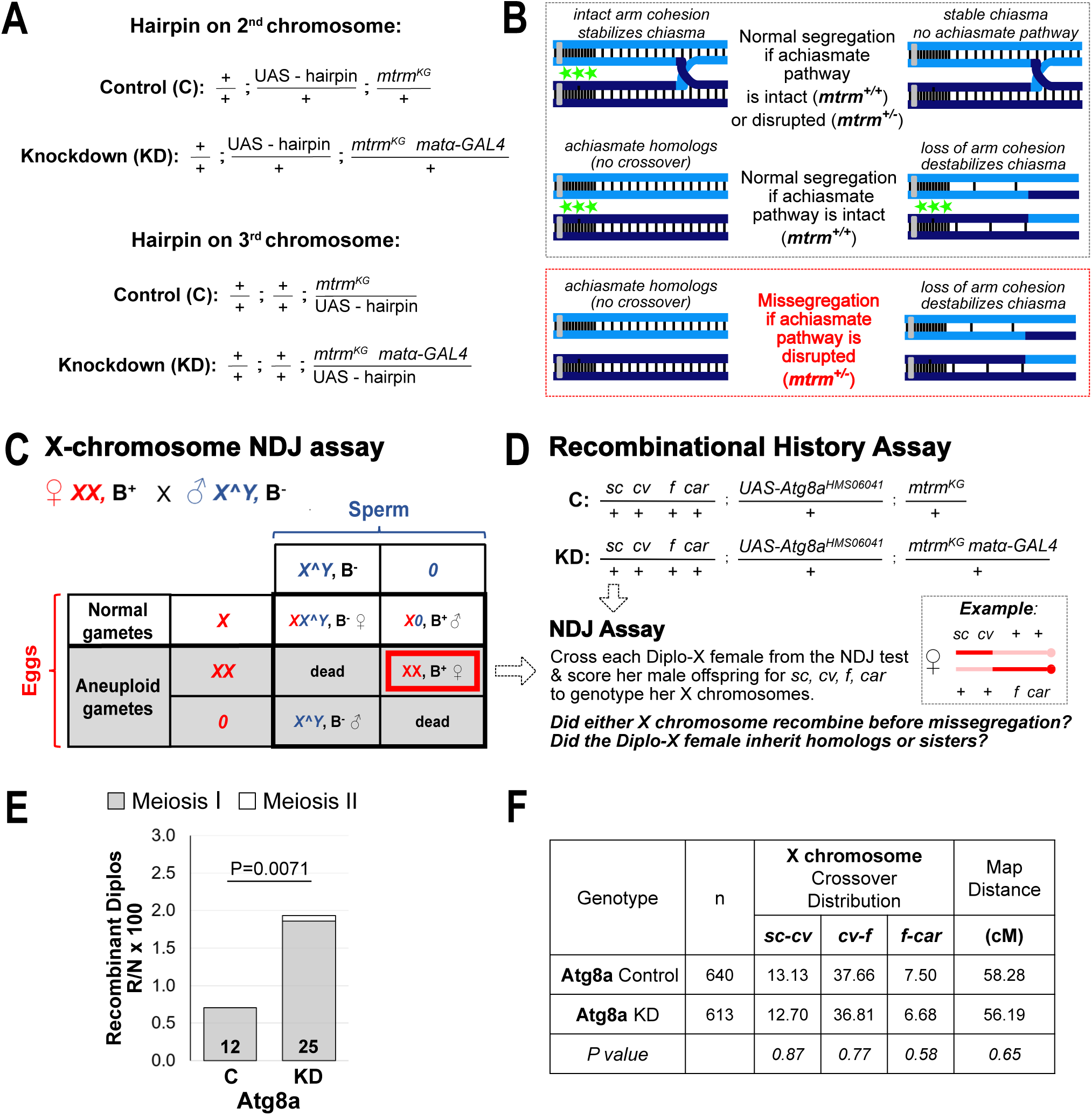
Missegregation in Atg8a KD oocytes is consistent with premature loss of cohesion. **(A)** Overview of genotypes used for NDJ tests. Flies are heterozygous for *mtrm^KG^* to disable the achiasmate segregation pathway in Drosophila oocytes. **(B)** The achiasmate system (depicted by green stars) relies on pericentric heterochromatin to keep bivalents that lack a crossover physically connected until anaphase I. However, this same system will also ensure accurate segregation of recombinant homologs for which premature loss of cohesion causes chiasma loss. In *mtrm^KG^/+* oocytes, premature loss of arm cohesion results in missegregation of recombinant bivalents. **(C)** We use the term nondisjunction (NDJ) to denote any type of missegregation event. In this NDJ test, virgins are crossed to *X^Y, B* males and their offspring scored for sex and eye phenotypes. Diplo-X female progeny (red box) inherit two X chromosomes from their mother because of missegregation. **(D)** By performing the original NDJ test with *sc cv f car/+* females, the X chromosome genotype of each Diplo-X female can be deduced by scoring her sons for the markers *sc, cv, f, car.* The centromere-proximal marker car allows one to distinguish between the missegregation of sisters or homologs. Arm cohesion, if maintained, should ensure that a recombinant bivalent remains associated and segregates accurately during anaphase I. Increased missegregation of recombinant homologs is consistent with premature loss of arm cohesion. (**E)** A NDJ test was performed with *Atg8a^HMS06041^*KD (matα driver) and control (no driver) females that were also *sc cv f car/+.* Diplo-X female progeny were genotyped using the Recombinational History Assay. Graph presents the frequency at which Diplo-X females inherited at least one recombinant chromosome. The frequency of missegregated homologs (Meiosis I) and sisters (Meiosis II) is plotted individually. The number of Diplo-X females scored is shown in the bar for each genotype. P-value shown is for meiosis I errors; meiosis II errors did not differ significantly. **(F)** Crossovers on the X chromosome were scored in Atg8a^HMS06041^ KD and control oocytes. Atg8a KD during meiotic prophase does not significantly impact the number or position of crossovers.

**Figure S3:**
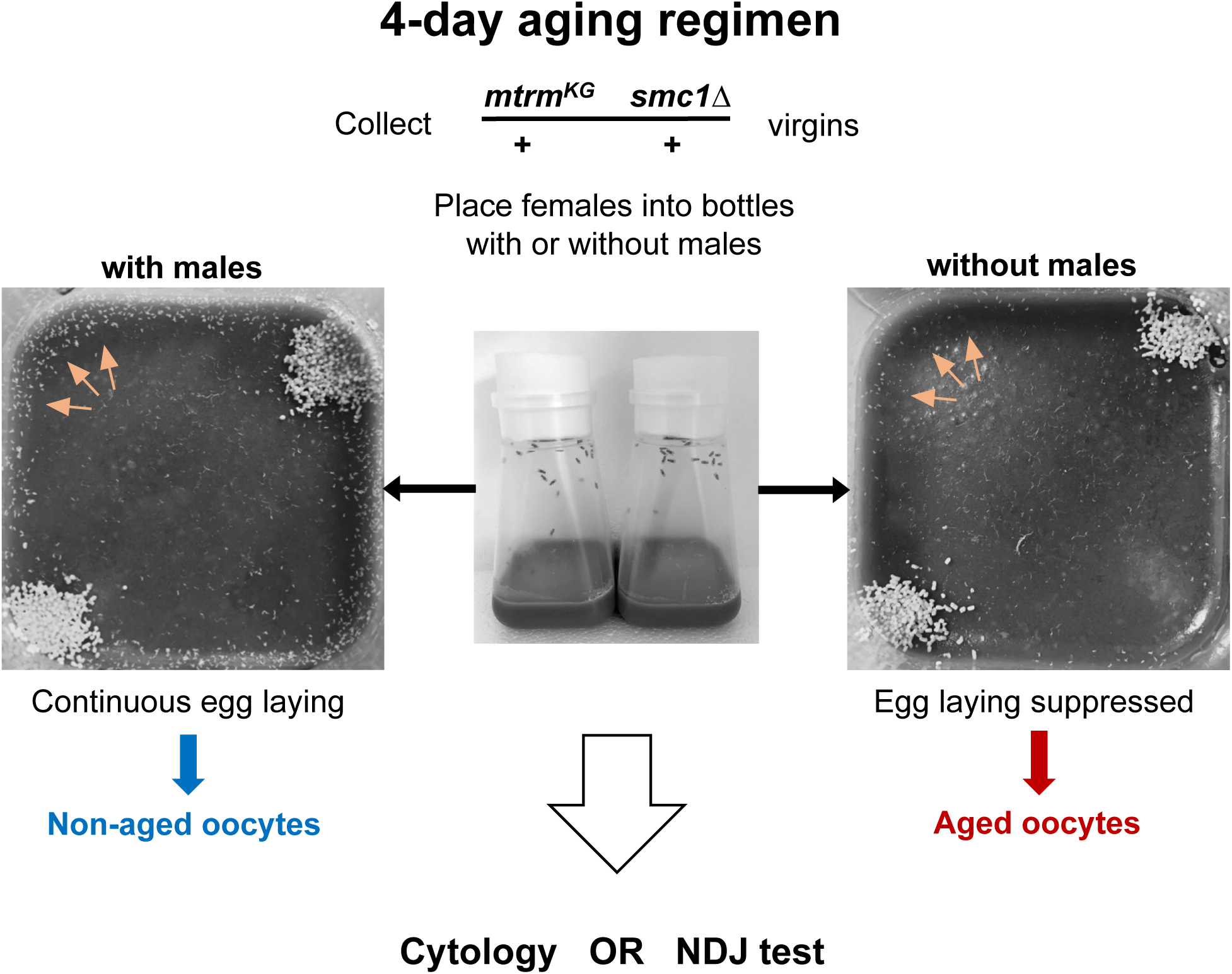
Procedure used to generate aged and non-aged oocytes under normal feeding conditions. In the past, for the 4-day aging regimen, we used a different food source (glucose-agar plates with a smear of wet yeast). This allowed easy replacement of the plate each day and photographic documentation of egg-laying or holding. However, given that diet affects autophagy and we are analyzing basal levels of autophagy, we modified our aging regimen to keep the food (cornmeal-molasses) constant throughout the entire experiment. Virgins (+/- males) are placed into normal fly bottles containing cornmeal-molasses based food with dry yeast sprinkled in two corners. **(Left)** Addition of *X^Y* males allows mating which stimulates egg laying and egg chambers move posteriorly through the ovariole as they grow. These females are the source of non-aged oocytes. **(Right)** If males are not added to the bottle, the virgins will not lay eggs and oogenesis halts, causing oocytes to “arrest and age” at different stages. These females are the source of aged oocytes. Flies are transferred to new bottles every 24 hours. Note that on the left, several eggs (embryos) have been laid on the food surface near the perimeter (arrows), but very few laid eggs (arrows) are visible on the right, verifying that virgin females are holding their eggs. Following the 4-day aging regimen, females can be mated to *X^Y* males to measure NDJ or ovaries can be dissected for cytology. *mtrm^KG^ smc1Δ / +* virgins were used for all aging experiments.

**Table S1:**
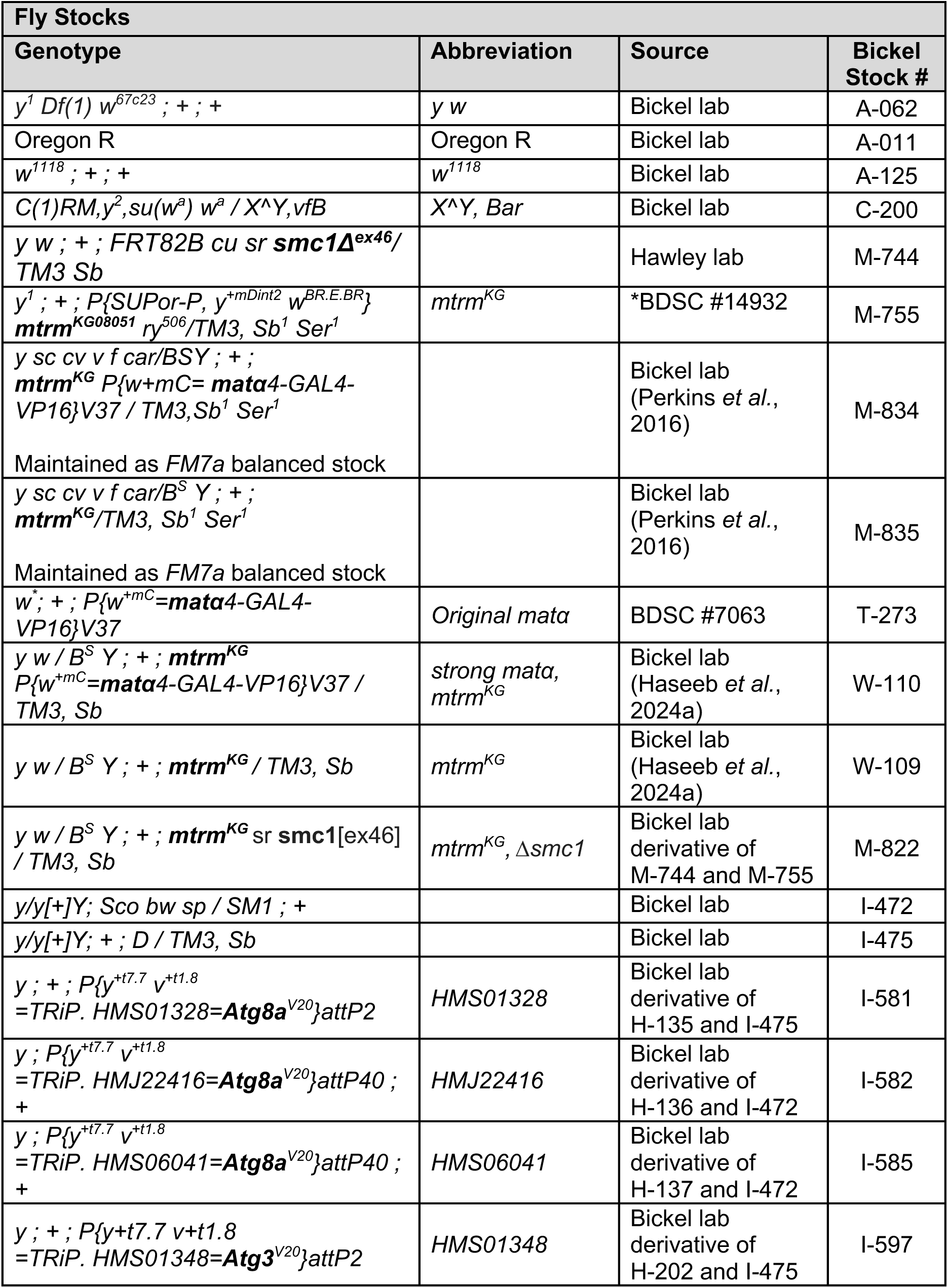
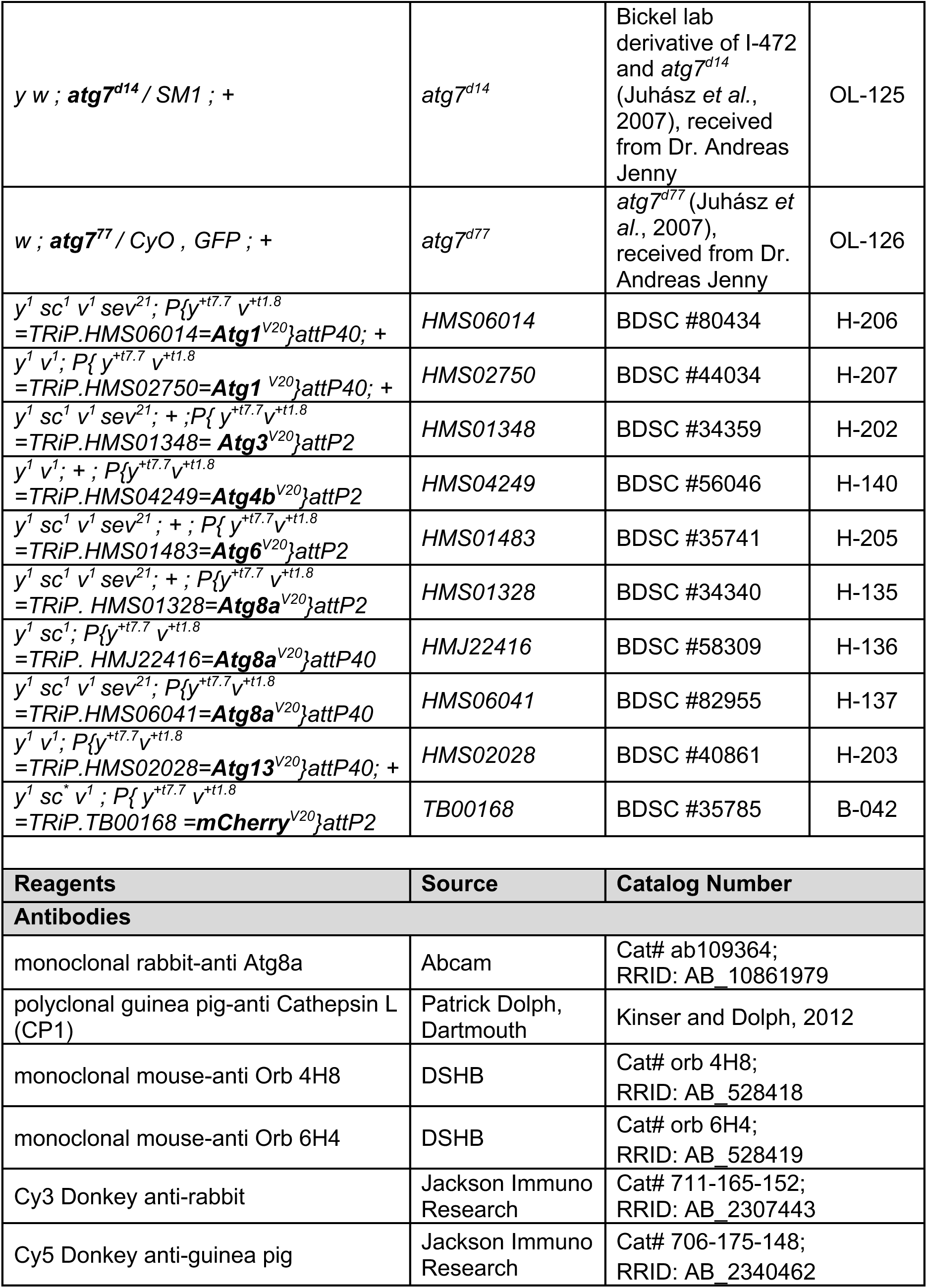
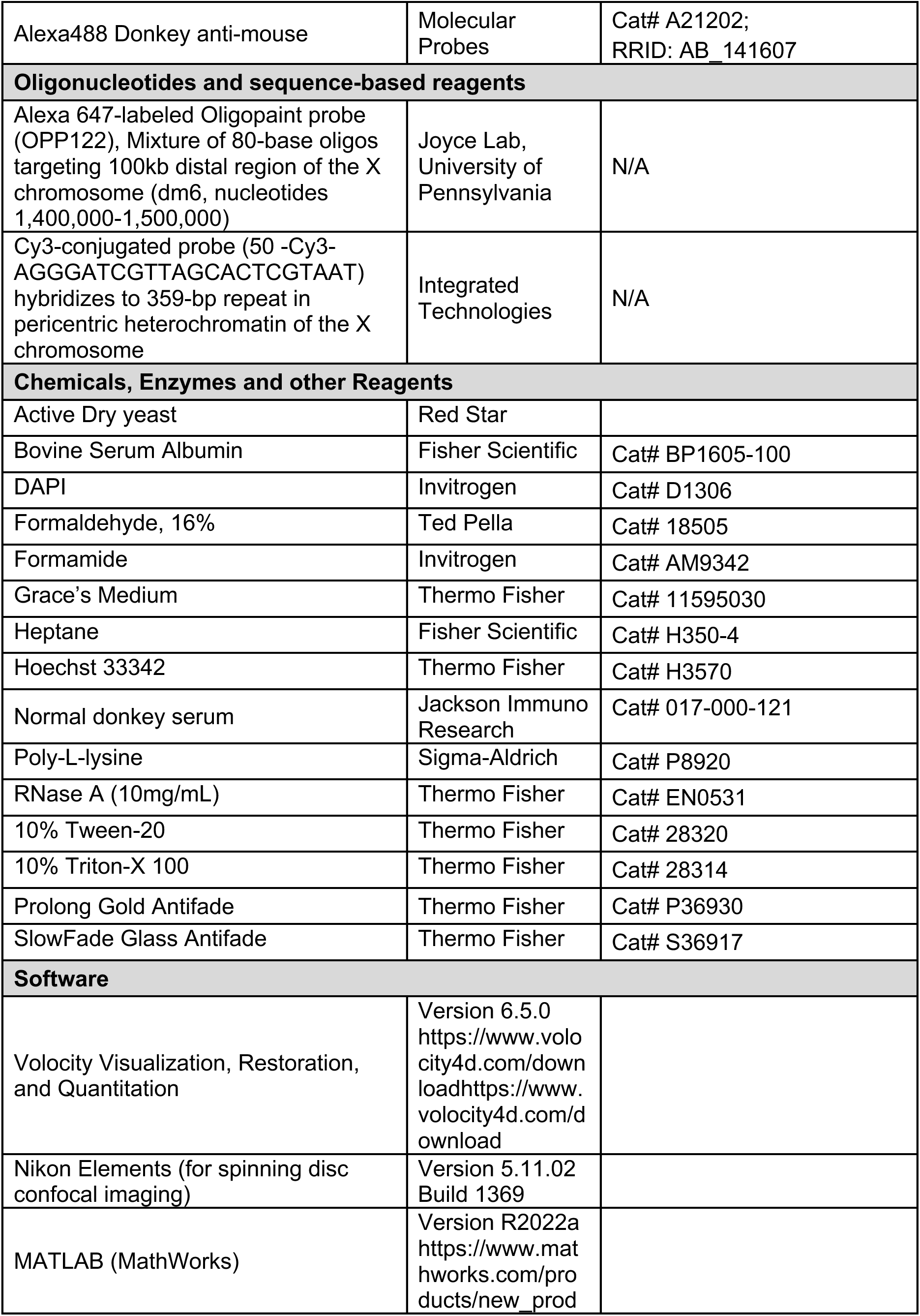
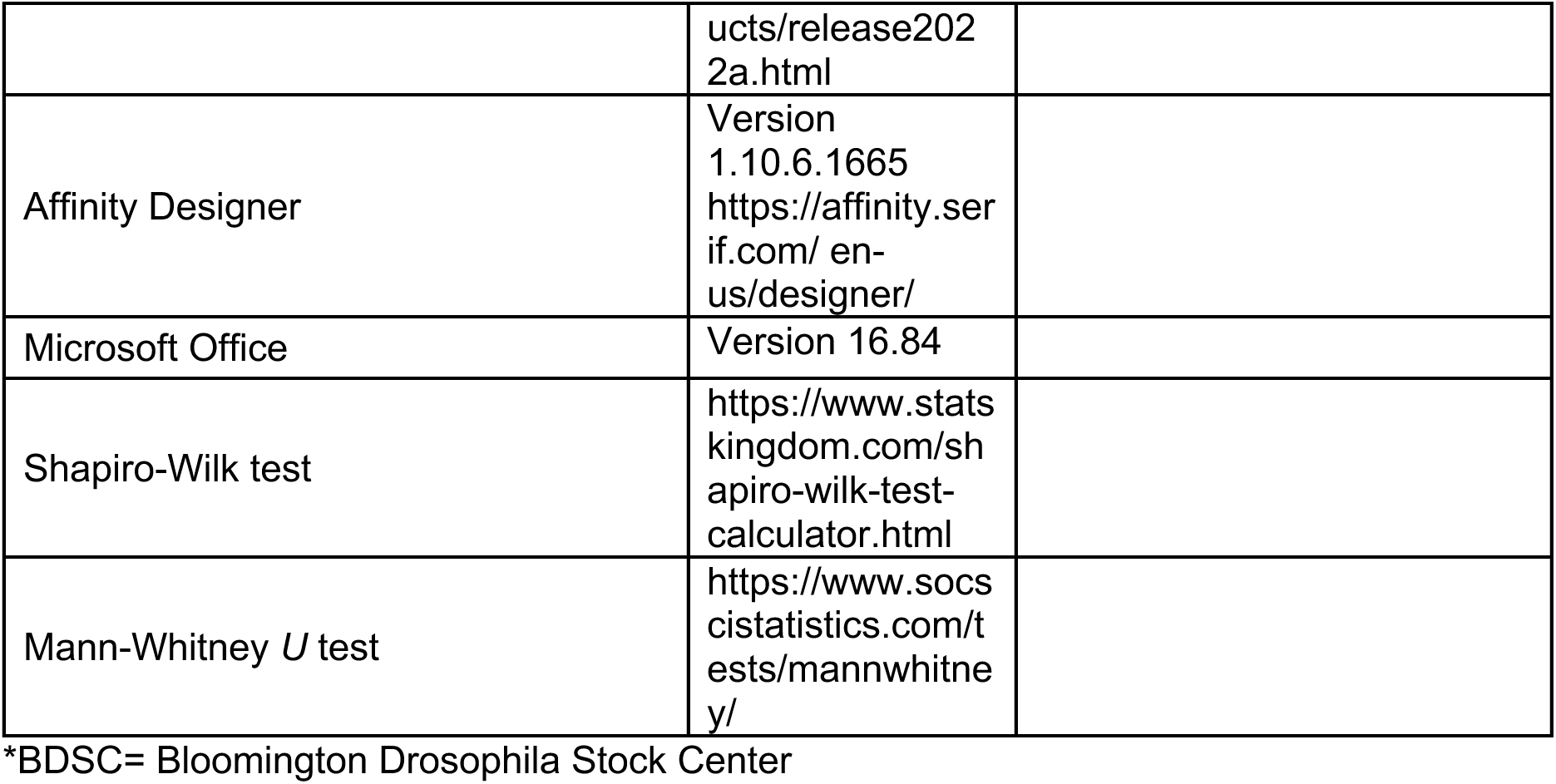
Fly Stocks and Reagents.

